# Northeast India: Genetic Inconsistency across Ethnicity and Geography

**DOI:** 10.1101/2024.01.09.574778

**Authors:** Biswabandhu Bankura, Bishnupriya Basak, Prajjval Pratap Singh, Albert Vanlalruata, Arindam Chatterjee, Sudakshina Ghosh, Rakesh Tamang, Manjil Hazarika, Gyaneshwer Chaubey, Madhusudan Das

**Affiliations:** Department of Zoology, University of Calcutta, Kolkata, India; Multidisciplinary Research Unit, Medical College Kolkata, Kolkata, India; Department of Archaeology, University of Calcutta, Kolkata, India; Cytogenetics Laboratory, Department of Zoology, Banaras Hindu University, Varanasi, India; Archaeology Wing, Art & Culture Department, Govt. of Mizoram, Aizawl, India; Department of Zoology, Vidyasagar College for Women, 39 Sankar Ghosh Ln, Kolkata-70006, India; Department of Archaeology, Cotton University, Guwahati, Assam, India

**Keywords:** Trans-Himalayan, Genetics, Migration, Ancestry, Mizo people

## Abstract

The genetic landscape of South Asia is intriguing. In a vast geographical region like India, mostly genes follow geography, whereas in a smaller geographical region e.g., Nepal, the genetic landscape is more akin to languages. Moreover, in neighboring regions like East and Southeast Asia, language is the major denominator of the genes. However, these are not just the two alternatives, in many geographical pockets, ethnicity plays a stronger role than either of these. Being a crossroads of South and Southeast Asia, the Himalayan region is mostly populated by people speaking the Trans-Himalayan language. Archaeological and anthropological studies suggest migration and cultural diffusion in this region with the Tibetan plateau in the North. To understand this complex pattern, we have performed a fine-grained genetic analysis of the major Mizo population and its various clans living in the Himalayan geography. We have investigated 110 individuals belonging to seven different clans of Mizo people for a hundred thousand autosomal markers. We have used various statistical methods and tested the role of ethnicity and geography in shaping the genetic landscape of the Himalayan region. In contrast to the East and Southeast Asian genetic landscape, our results suggested that the fine-scaled genetic structure of Northeast India is much more complex and a mixture of interplay between languages, ethnicity, and geography. Allelic frequency-based analyses indicated that a novel Trans-Himalayan ancestry unite all the populations of this region.

## Introduction

The eastern Himalayan region, extending from Sikkim in the West to Arunachal Pradesh in the East, played a very crucial role in shaping the population history of the Indian subcontinent. This area constitutes a unique narrow passageway that connects the Indian subcontinent with East Asia and Southeast Asia. The archaeological evidence, although fragmentary and meagre, suggest population movements and cultural connections on both sides of the Himalayan region, particularly with the Tibetan plateau in the North [1].

The Northeastern region of India consists of the states of Arunachal Pradesh, Assam, Manipur, Meghalaya, Mizoram, Nagaland, Sikkim, and Tripura. Mizoram is the southernmost state in the northeast, sharing its borders with Tripura, Assam, and Manipur along with Myanmar and Bangladesh. The Mizo (a.k.a. Lushai) people are believed to be a part of the Trans-Himalayan stock. The Mizos linguistically belong to the Kukish (a.k.a. Mizo-Kuki-Chin) group of the Tibeto-Burman (now Trans-Himalayan) language family[2]. Mizoram has the highest number of tribal people among all states of India[3]. The majority of Mizoram’s population consists of several ethnic tribes who are linked with each other historically, culturally, or linguistically. These ethnic groups are collectively known as “Mizos” (Supplementary Text).

*Mizo* means ‘people of the hills’. “Mizo”, is a constructed concept with a foundation in collective memory and myth (Supplementary Text). There are several theories regarding the migration of Mizos to the present state of Mizoram. Most of our understanding of the early history of the Mizos is based on traditions, folklore, beliefs, legends, and customs. One of the most accepted theories is that the ancestors of Mizos had their earliest settlement in south-eastern China, in a place or mythological cave called Chhinlung[4]. The linguistic affinity, mode of living, and culture of the Mizos with hill tribes of southern China provide some clues for their common origin, place, and ancestors, due to which southern China can be considered as the original homeland for the Mizos[5]. From Chhinlung, they moved southward to the Shan state of Myanmar, then to Kabaw Valley and the Chin Hills subsequently. This theory of the origin and migration is common among most sub-tribes of the Mizos[6],[7],[8].

Genetic simulation data strongly suggest that the Trans-Himalaya region, harboured human populations and even must have acted as a thoroughfare for human migration since the Early Upper Palaeolithic[9],[10],[11],[12],[13]. Yet the precise role of this region in the peopling of Eurasia remains only poorly understood, and several prominent questions remain unanswered with regard to the appearance and dispersal of modern humans in this region. Located at the nexus of the hypothetical Southern vs. Northern route dispersals, Trans-Himalaya represents a huge lacuna in the genetic data. In the Trans-Himalayan region previous studies done on lower resolution and (or) smaller number of samples, suggested genetic discontinuity between Northeast India and mainland India[14].

Therefore, in this study, we have generated new Genome-wide data on more than 100 samples belonging to various clans of the Mizo population and reconstructed the population history of the Trans-Himalayan region with a particular focus on the population history of Mizo. We generated the genome-wide data of nearly half a million SNP and merged it with 1082 samples from Eurasian populations. We took advantage of the large samples and high-resolution data, and with fine-scaled genetic analysis, we tested for the factors playing roles in shaping the genetic landscape of the Trans-Himalayan region. Research into fine-scale genetic variation is rapidly advancing, and it is expected to continue to provide important insights into the evolutionary history of human populations as well as the genetic basis of health and diseases.

## Results and Discussions

The large number of samples incorporated in this study helped us to elucidate the fine-scaled genetic variation of the Trans-Himalayan region and their relation with other Eurasian populations. For the first time, we used a haplotype-based analysis to detect the subtle level of genetic variation (Fig. 1). We used the Mizo population as a model which has a distinct clan structure (Supplementary Figure 1). At the Eurasian level of population differentiation, the Trans-Himalayan populations form a related but distinct cluster with the East and Southeast Asian populations (Supplementary Figure 2). Mizo also occupies the same clade as Trans-Himalayan, suggesting its origin from the same stock population.

**Fig. 1:**
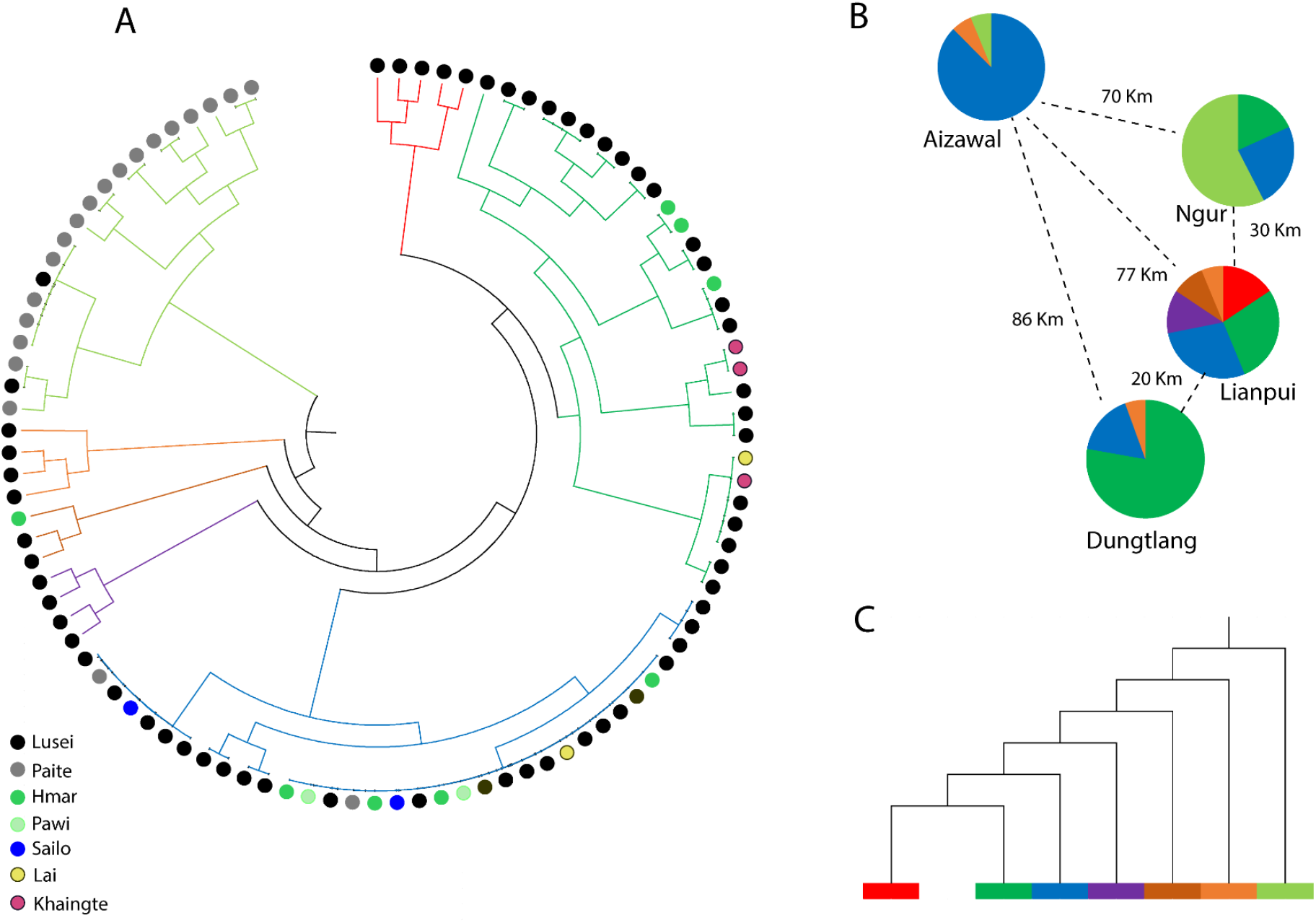
A) The Maximum Likelihood (ML) subtree showing the clan wise clustering of Mizo. B) The district wise cluster proportions and their arial distances from one another. C) The phylogenetic relation of the seven clusters emerged from fineStructure analysis.

At the micro level population differentiation, our analysis detected seven genetic clusters among the Mizo clans (Fig. 1A). With some exclusive occupancy, most of the clans are distributed among more than one cluster. We divided these individuals based on their geographic locations (districts) and asked which factor shaped the present population differentiation. Interestingly, even at the smaller geographical distances, we see a distinct population’s genetic structure (Fig. 1B). Moreover, this population’s genetic structure did not follow pure phylogenetic structure (Fig. 1C). Thus, the population structure of Trans-Himalayan is much more complex. It cannot be explained solely based on language, geography, or ethnicity. It is rather an outcome of the interplay between all these factors.

At the Eurasian population comparison, the Mizo formed a close cluster overlapping with the East Asian populations (Fig. 2). However, it formed a different cline as the East Asian population. It followed the same cline as other Trans-Himalayan populations[9]. This cline is different from Southeast Asian cline and placed in between East and South Asia, and exclusively carried the Trans-Himalayan populations. The ADMIXTURE analysis supported previous work[9] suggesting the presence of a major Trans-Himalayan genetic component. Mizo population has a unique ancestry which is also shared by other Indian Trans-Himalayan populations. Dark green colour and light green colour represent South Asian specific component while the orange and dark pink colour represents East Asian and Mizo-specific component respectively. Likewise, the outgroup *f3* statistics suggested a closer genetic affinity of Mizo with the Trans-Himalayan populations (Fig. 4).

**Fig. 2:**
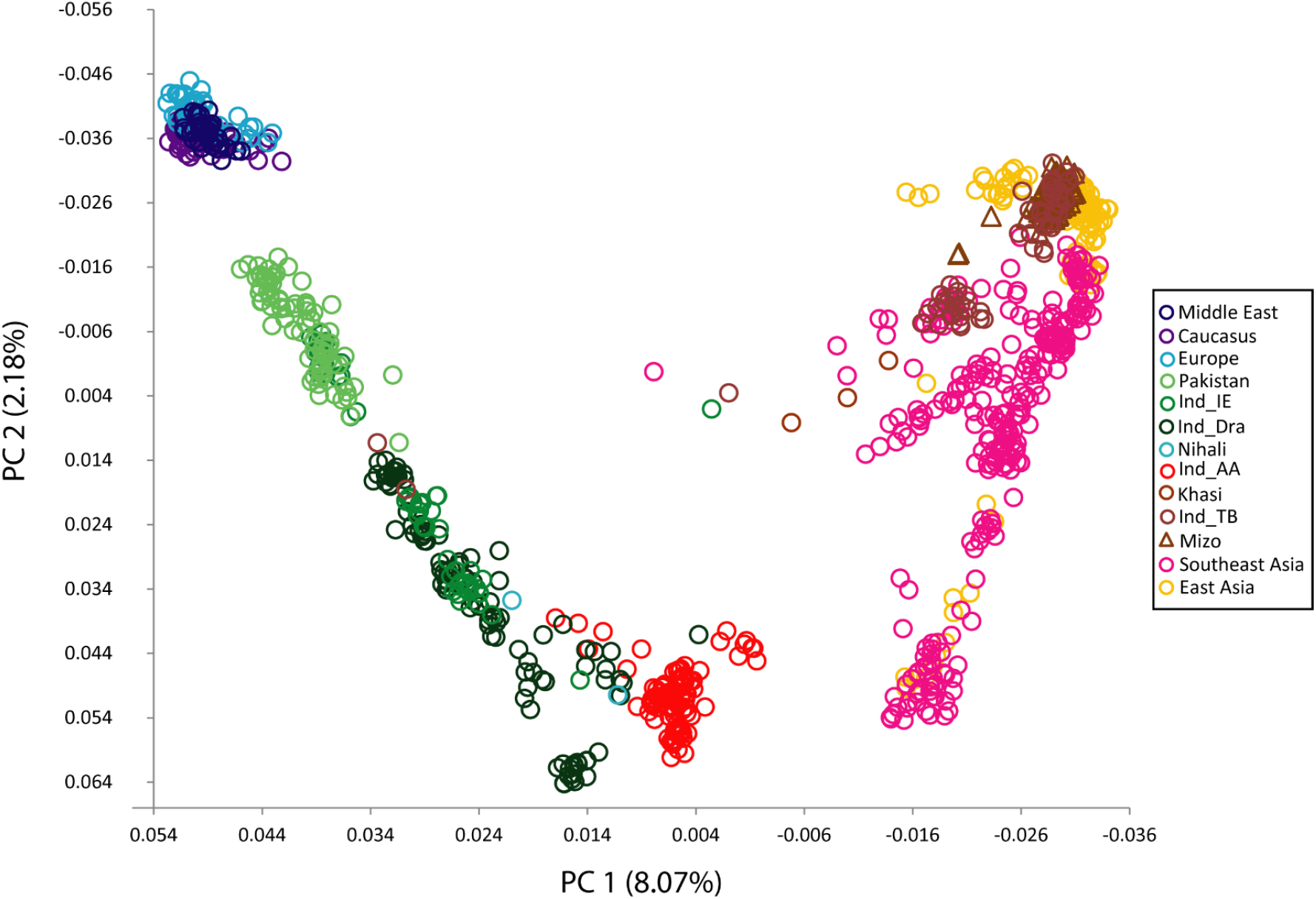
The Principal component (PC) analysis of Eurasian population showing the placement of Mizo cluster.

**Fig. 3:**
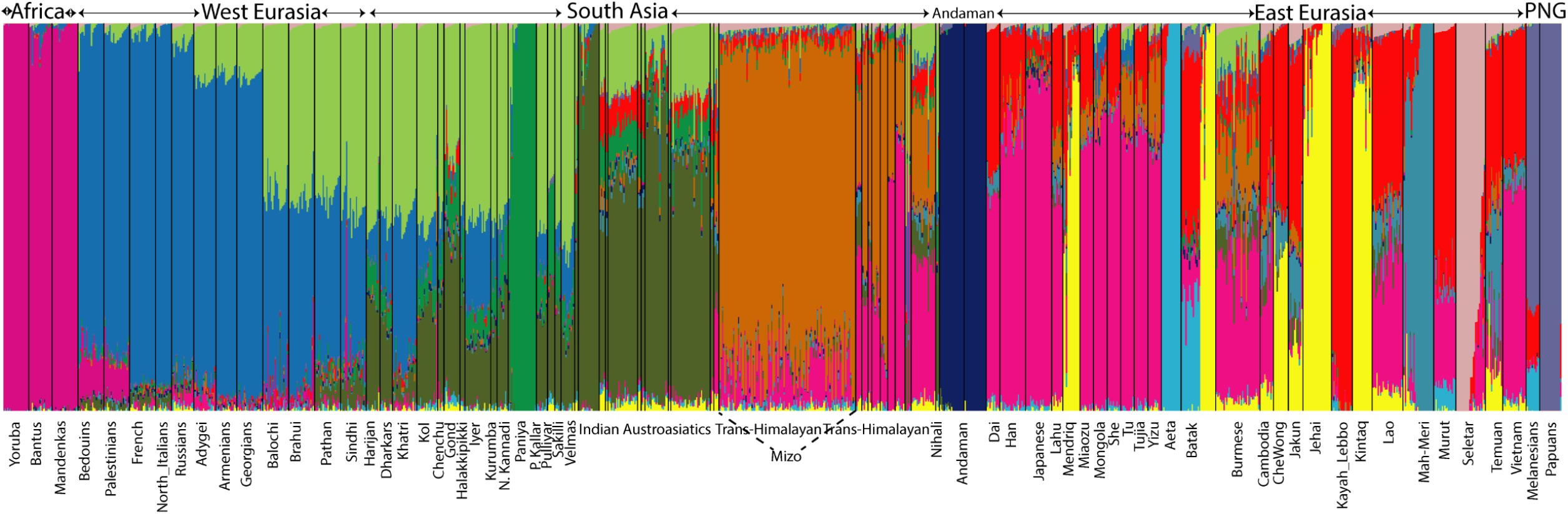
The ADMIXTURE analysis showing the component sharing of Mizo and Trans-Himalayan populations.

**Fig. 4:**
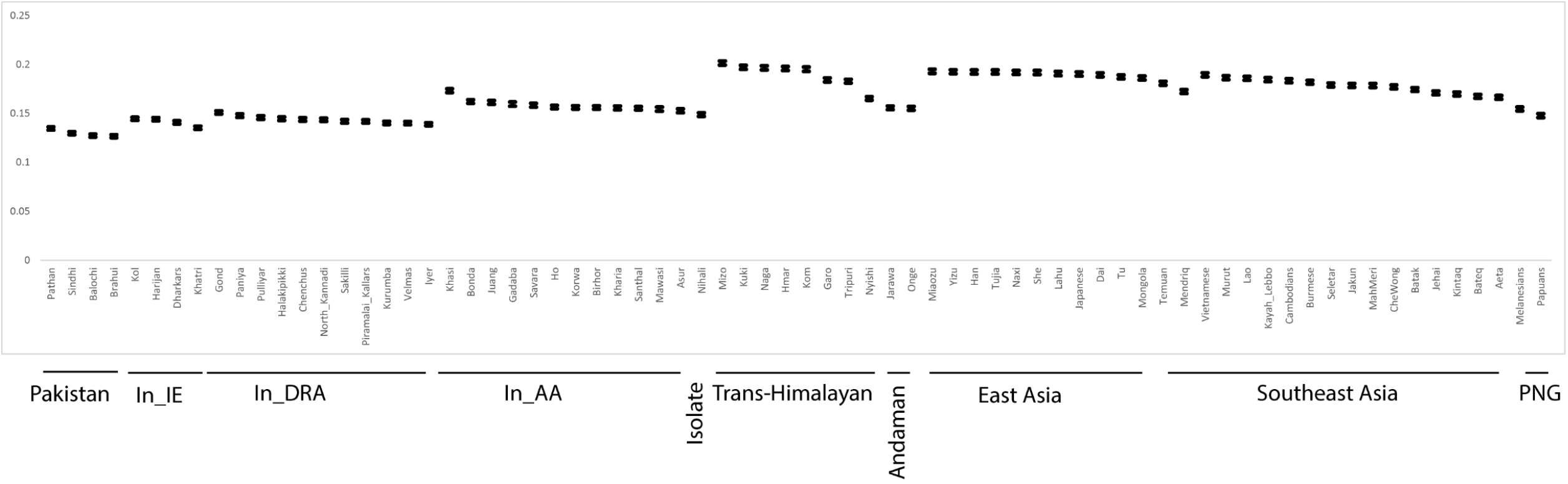
The outgroup *f3* statistics of Mizo showing the highest shared drift with the Trans-Himalayan populations.

The Runs of Homozygosity (RoH) yielded variable results for various clans (Fig. 5), suggesting genetic heterogeneity in the clan marriage pattern. The time of admixture for the East and South Asian ancestries for the Mizo population has yielded a timeline of 1650-2050 years suggesting a later migration than the Austroasiatic Munda populations[15]. This date may pick one of the signals of Trans-Himalayan expansion to South Asia which has been thought to be multilayered[10].

**Fig. 5:**
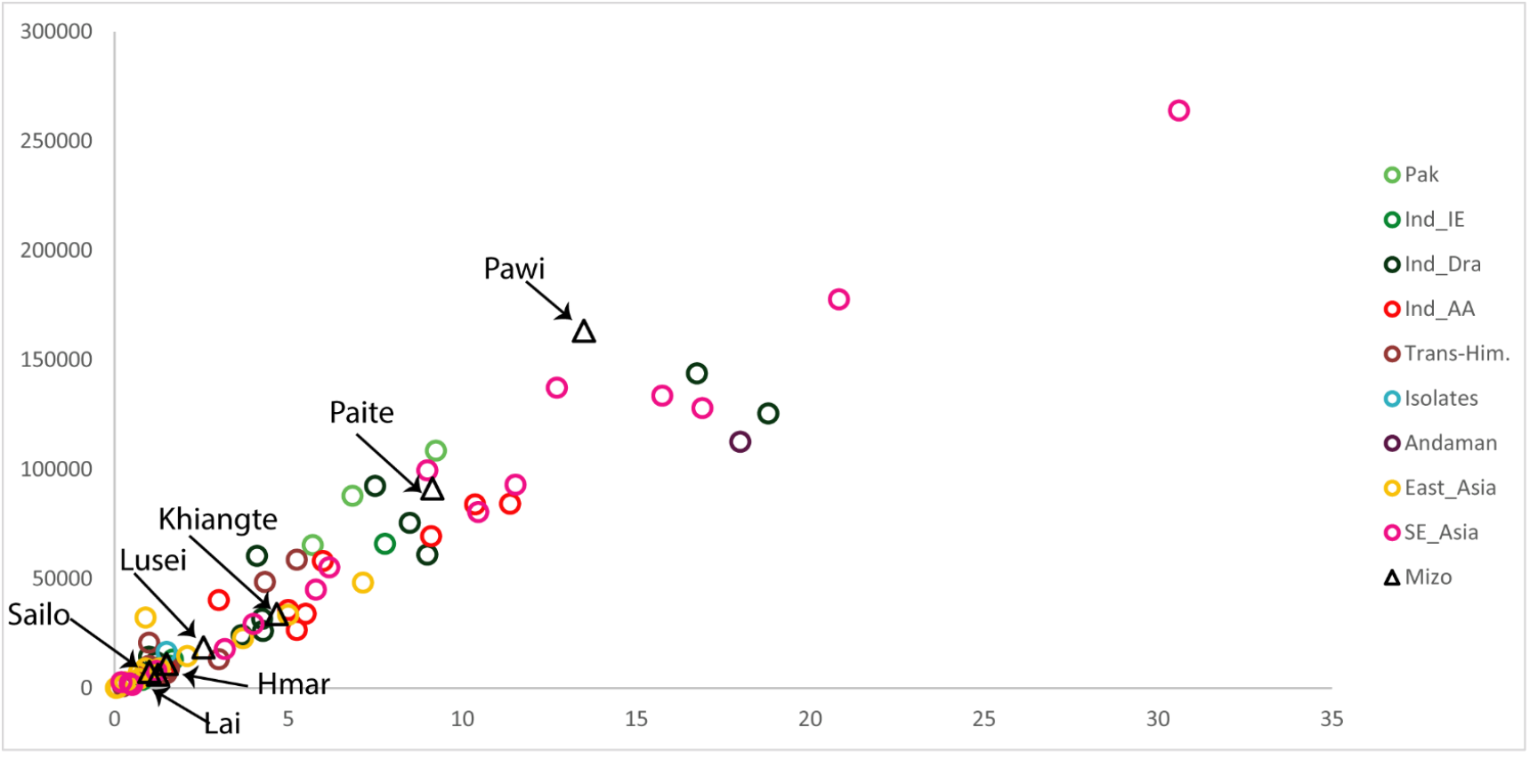
The Runs of Homozygosity (RoH) analysis of various clans of Mizo.

In conclusion, our extensive fine-grained analyses establish a strong point on the genetic history of the Mizo people that they are a distinct community of South Asia that is genetically closer to East Asian specific ancestry, but they are quite differentiated from them and belong to the same stock of other Indian Trans-Himalayan people. The findings of the study suggested a complex genetic structure in Northeast India, contrasting with the genetic landscape observed in East and Southeast Asia. The results indicated that the genetic makeup of the Mizo population and its clans in the Himalayan region is influenced by a mixture of languages, ethnicity, and geography. Overall, this study sheds light on the intricate genetic patterns in the Himalayan region, emphasizing the role of languages, ethnicity, and geography in shaping the genetic landscape of Northeast India.

## Materials and Methods

This study is approved by the Ethical Committee of the University of Calcutta. Sampling was executed in the border area and we have eluded up to three generations for sampling (Supplementary Figure 1). Blood samples were collected from healthy individuals belonging to the Mizo population with informed consent. DNA was isolated from blood following the standard protocol[15], and was sequenced using Illumina global sequencing array platform. Obtained data were processed/curated by using PLINK 2.0[16] with the --mind -–geno -–maf command. We merged our newly generated dataset to answer our questions with available world data published by the Institute of Genomics, Estonia[17]. We pruned merged data to eliminate background linkage disequilibrium (LD) by eliminating one SNP of any pair in strong LD r^2^ > 0.4, in a window of 200 SNPs (set the sliding the window by 25 SNPs at a time) that can affect both principal component analysis (PCA) and ADMIXTURE analysis. Data were classified based on the location while Indian populations were classified based on languages such as Indo-European, Austro-Asiatic, Dravidian, and Tibeto-Burman. EIGENSOFT[18] package was used to perform PC analysis for analyzing the population structure and genetic variation with the default setting. Unpruned data was used for other analyses except for ADMIXTURE and PCA. ADMIXTURE[19] was run for visualizing the ancestral component sharing, for our merged dataset the best supported ancestral component (K) was K=14. In order to understand a more detailed population structure, fineSTRUCTURE ver 1.0[20] was run and Chromopainter[20] was performed to paint the chromosome of donors and recipients. BEAGLE 5[21] was used to phase the data set for finSTRUCTURE and Chromopainter tool. MEGA version 7[22] was used to finalize the Maximum likelihood tree obtained from fineSTRUCTURE analysis. Different F statistics from ADMIXTOOL2.0[23] package were performed to know the direction of gene flow and to calculate the shared drift. ALDER 1.0[24] was used with the default setting to know the admixture timing with their source populations. Runs of homozygosity (RoH)[25] were used to understand the demographic past of the Mizo population.

## Acknowledgment

The authors would like to thank Mr. Pachuau Rohmingthanga, Mr. Singson Mahuma, Mr. Mapuia Hnamte, District Administration Champhai, Department of History & Ethnography Mizoram University, and, Prof. Samir Das (Department of Political Science, University of Calcutta) for their kind help in this research work. This work was supported by University Grant Commission/ University with Potential for Excellence II program [Ref. No.: UGC/521/UPE-II/Look East, Date-12-07-2017].

## Author Contribution

The conception of the study, reviewing the draft, Study design: M.D; Literature search, data collection, experimental design, result analysis, making draft: B.B, B.B; Literature Search, Statistics, result analysis, making draft: P.P.S; data collection: A.V; experimental design: A.C; Literature search, Review Draft: S.D; Literature search, Review Draft: R.K; Literature search, making draft, Review Draft: M.H; Statistics, making the draft, Literature Search, reviewing the draft, Interpretation & Analysis: G.C

## Declaration of interests

The authors declare that they have no competing interests.

## Notes

### Competing Interest Statement

The authors have declared no competing interest.

